# Cardiac metabolic imaging using hyperpolarized [1-13C]lactate as a substrate

**DOI:** 10.1101/2020.12.04.412296

**Authors:** Angus Z Lau, Albert P Chen, Charles H Cunningham

## Abstract

Hyperpolarized [1-^13^C]lactate is an attractive alternative to [1-13C]pyruvate as a substrate to investigate cardiac metabolism in vivo; it can be administered safely at a higher dose and can be polarized to a similar degree as pyruvate via dynamic nuclear polarization. While ^13^C cardiac experiments using HP lactate have been performed in small animal models, it has not been demonstrated in large animal models or humans. Utilizing the same hardware and data acquisition methods used in the first human HP ^13^C cardiac study, 13C metabolic images were acquired following injections of HP [1-^13^C]lactate in porcine hearts. Data were also acquired using HP [1-^13^C]pyruvate for comparison. The ^13^C bicarbonate signal was localized to the myocardium and had a similar appearance with both substrates for all animals. No ^13^C pyruvate signal was detected in the experiments following injection of hyperpolarized ^13^C lactate. The SNR of injected lactate was 88 +/-14% of the SNR of injected pyruvate, and the SNR of bicarbonate in the experiments using lactate as the substrate was 52+/-19% of the SNR in the experiments using pyruvate as the substrate. The lower SNR was likely due to the shorter T1 of [1-^13^C]lactate as compared to [1-^13^C]pyruvate and the additional enzyme-catalyzed metabolic conversion step before the 13C nuclei from [1-13C]lactate were detected as ^13^C bicarbonate. While challenges remain, the potential of HP lactate as a substrate for clinical metabolic imaging of human heart was demonstrated.

## Introduction

Hyperpolarized (HP) ^13^C MRI is a promising tool for non-invasive characterization of *in vivo* metabolism (1,2). It has been demonstrated in small and large animal models that HP MRI can be used to probe changes in cardiac metabolism due to modulation in substrate availability as well as cardiac diseases (3-5). The feasibility of studying cardiac metabolism in the human heart has also been shown recently (6,7). In all the hyperpolarized metabolic imaging studies performed so far, [1-^13^C]pyruvate is the most commonly used substrate due to the high achievable polarization, long T_1_ relaxation time, and the important role of pyruvate in cellular metabolism. However, pyruvate is typically injected in supraphysiological concentrations (> 0.1 mmol/kg), potentially altering the metabolic system that is being interrogated. [1-^13^C]lactate is an attractive alternative; it can be given safely at high millimolar physiological concentrations and can be polarized to a similar degree as pyruvate via dynamic nuclear polarization (DNP) (8-11). Potentially, PDH flux can be interrogated with less disturbance of the metabolic state by measuring HP pyruvate and bicarbonate signal within the cardiac muscle produced from the HP lactate substrate (9).

While ^13^C cardiac experiments using HP lactate have been performed in small animal models by various groups (8-11), it has not been demonstrated in large animal models or humans. Utilizing the same hardware and data acquisition methods that have been used in the initial human HP ^13^C cardiac study (6), we investigated the use of hyperpolarized [1-^13^C]lactic acid as a probe of cardiac metabolism in pigs as a large animal model of the human heart. Data were acquired using HP [1-^13^C]pyruvate as the substrate in the same animals for comparison.

## Methods

### Hardware and sample preparation

All scans were performed using a 3T MR system (GE MR750, GE Healthcare, Waukesha, MI). A ^13^C “clamshell” volume transmit coil and an 8 channel receive coil system that consist of 2 separate paddles containing 4 receive elements each were used (GE). The pigs were placed supine and feet first inside the transmit coil, with one of the receive paddle array placed on the anterior chest wall over the heart. [1-^13^C]lactic acid (with 20% water) and neat [1-^13^C]pyruvic acid (Isotec, Miamisburg, OH) were mixed with 15 mM trityl radical (GE) and polarized in a GE SpinLab DNP polarizer. Details on the lactic acid sample preparation are included in supplementary material. 250 mg of either substrate was loaded into the sample vial, but since the lactic acid sample is less concentrated, an additional amount of water was added in the receiver for the pyruvate sample (in addition to the TRIS/NaOH neutralization solution) to achieve the same final concentration for both substrates. Following dissolution and neutralization, the substrate (either lactate or pyruvate) at 125 mM in 15 mL was injected at a rate of 1 mL/s, followed by 5 mL saline flush.

### ^13^C cardiac MRS/MRI protocols

All animal experiments were carried out under a protocol approved by the institutional animal care and use committee. Animal preparation and handling procedures followed the same methods described previously (12). Three Yorkshire pigs were used in these experiments (30-33 kg). Three infusions of HP substrates (2 lactate, 1 pyruvate) were performed during each experiment, approximately 30 minutes apart.

For the dynamic MRS experiments, a pulse-acquire pulse sequence with 9 degree flip angle and a cardiac gated repetition time of ∼2 s (3 or 4 R-R intervals) was used. The data acquisition started at the same time as the injection and 64 temporally resolved spectra were acquired. For the ^13^C imaging experiments, short-axis images of ^13^C-bicarbonate, [1-^13^C]lactate, and [1-^13^C]pyruvate were obtained using a multi-slice, spectrally-selective spiral imaging pulse sequence covering the left ventricle as previously used in both pre-clinical large animal models and humans studies (1×1×1 cm^3^ resolution, 6 slices, scan duration 18 cardiac cycles) (6,12). Data acquisition started following a delay from the time of injection determined from the dynamic MRS data from each subject. The experimental protocol is summarized in Table 1. For experiments using lactate, the metabolite acquisition order was modified to preserve the magnetization of the substrate (Figure 1). The_spiral images were reconstructed by non-uniform fast Fourier transform. A 10 Hz exponential filter was applied to the time-domain spiral data, resulting in a final in-plane resolution of 13×13 mm^2^. Images from the individual coil channels were combined using a root sum-of-squares combination. The maximum SNR for each metabolite image was calculated by dividing the maximum signal by the standard deviation over the image background.

**Table 1.**
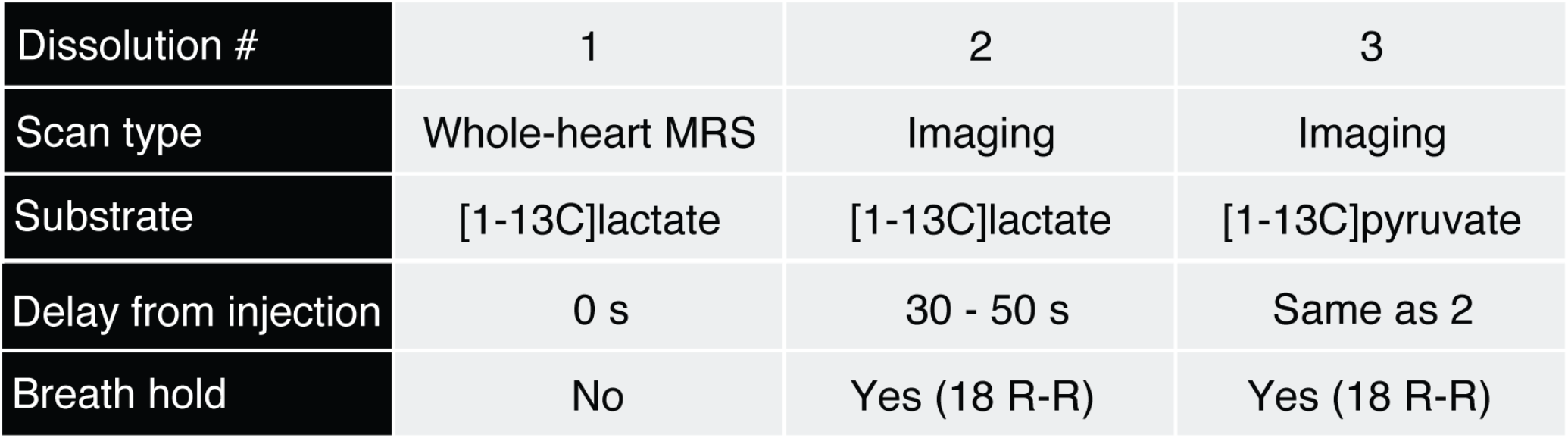
Summary of MR experiments performed on each subject

**Figure 1.**
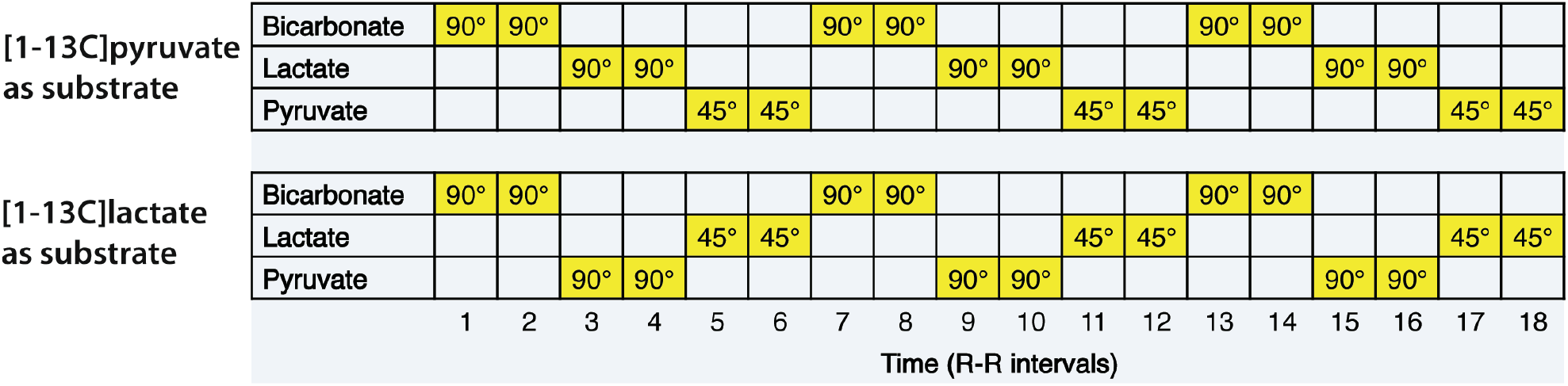
Flip angle schedule of the multi-slice spectrally-selective spiral imaging pulse sequence used for 13C cardiac imaging following injection of HP [1-13C]pyruvate or [1-13C]lactate.

## Results

Representative dynamic ^13^C spectra acquired from pig heart *in vivo* following injection of HP [1-^13^C]lactate are shown in Fig. 2. In addition to the substrate resonance, [1-^13^C]pyruvate, [1-^13^C]alanine and ^13^C bicarbonate resonances were also observed. The first 30 spectra from the dynamic MRS data (Fig. 2 left) as well as the spectrum from the time point with maximum ^13^C bicarbonate signal (Fig. 2 right) are shown. The time courses of the HP [1-^13^C]lactate and ^13^C bicarbonate signals for the three animals are plotted in Fig 3. Note the shape of the substrate bolus as well as the time from injection to temporal maximum for both substrate and metabolites were somewhat different for the three subjects. This was likely due to variability associated with the manual injections. Physiological differences between the subjects may also play a role. While the injected lactate time-to-peak from the start of injection had a larger range of values at 19 s, 13 s, and 25 s, the lactate and bicarbonate peak-to-peak duration were more similar at 19 s, 24 s and 20 s. The imaging window for the subsequent 13C MR imaging experiments were chosen so the start of the imaging acquisition was at the temporal maximum of the 13C bicarbonate (shown in Fig. 3 as green blocks).

**Figure 2.**
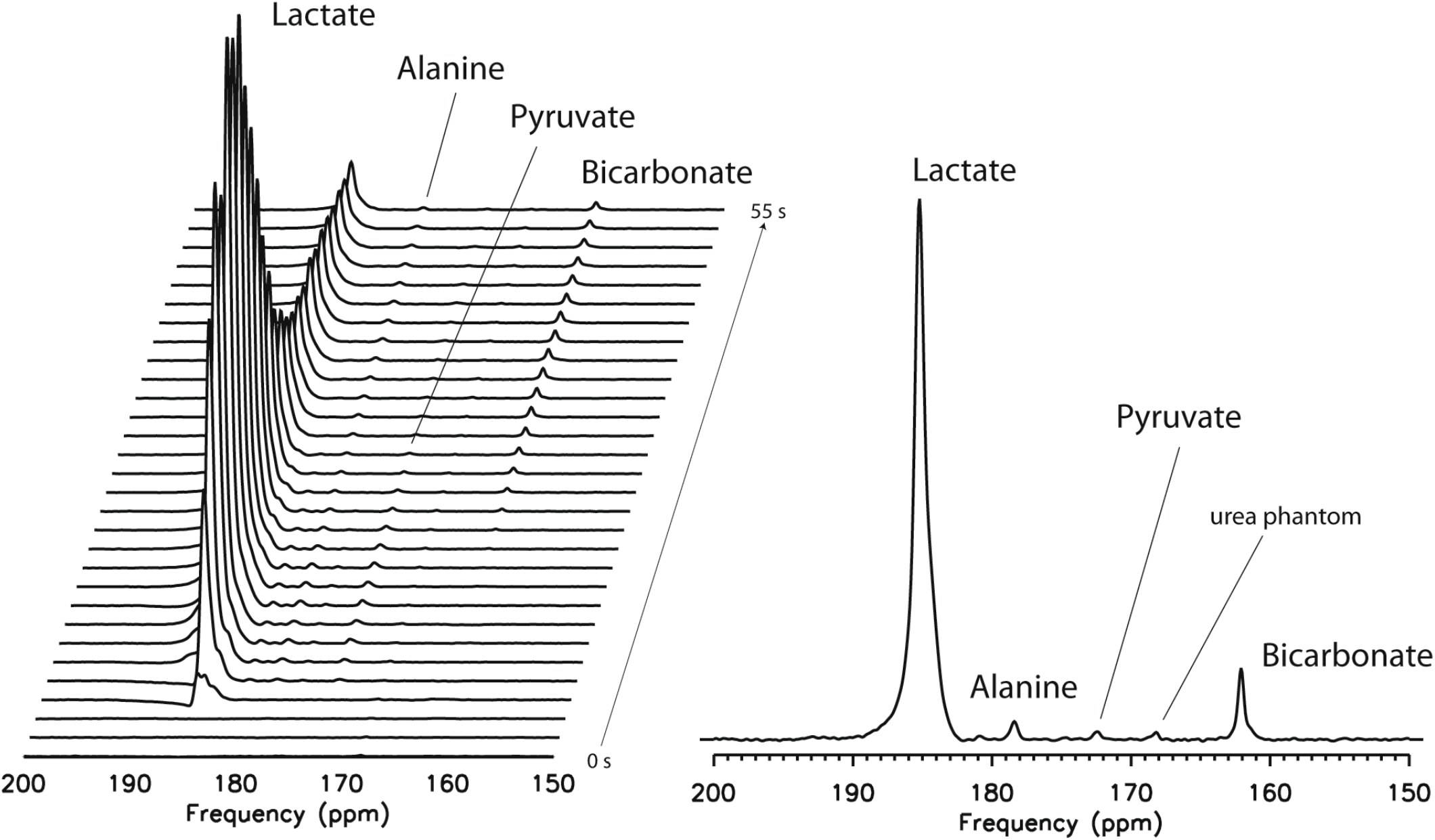
Representative 13C spectra acquired from a pig heart in vivo following injection of HP [1-13C]lactate. The first 30 spectra from the dynamic MRS dataset (left) as well as the spectrum with the maximum 13C bicarbonate signal (right) are shown.

**Figure 3.**
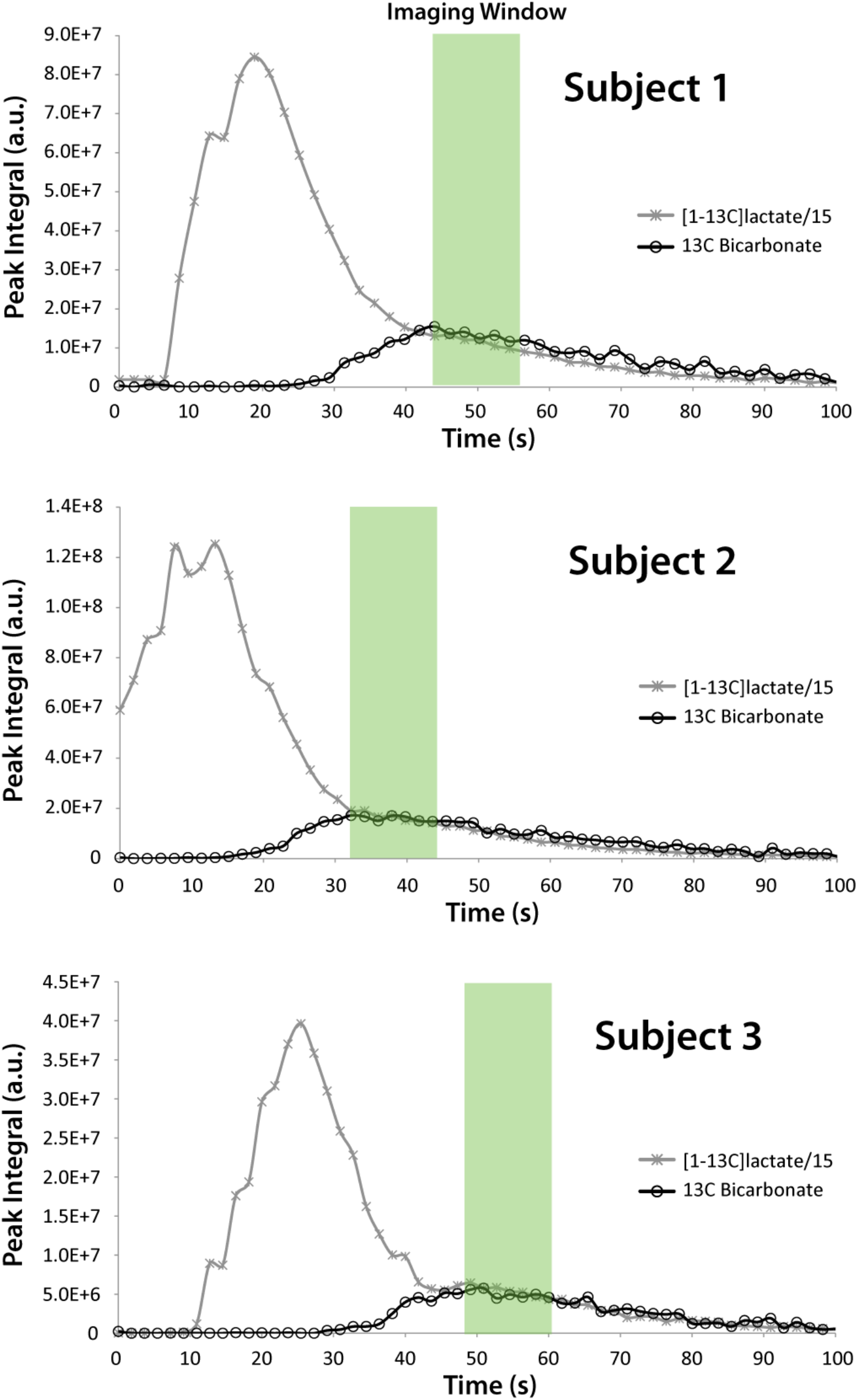
Signal time course for the injected HP [1-13C]lactate substrate and 13C bicarbonate obtained from the dynamic MRS experiments. The imaging windows for the subsequent 13C imaging experiments are indicated by the green blocks.

Images of hyperpolarized ^13^C lactate, pyruvate and bicarbonate from the hearts of the three animals are shown in Fig. 4. The ^13^C substrate signals, whether it was lactate or pyruvate, were localized primarily in the cardiac chambers. The ^13^C bicarbonate signals were localized in the myocardium and had a similar appearance using either substrate for all animals. No ^13^C pyruvate signals were detected in the experiments following injection of hyperpolarized ^13^C lactate. ^13^C lactate images obtained in the experiments following injection of hyperpolarized ^13^C pyruvate showed a more diffuse spatial distribution, similar to data observed previously (12). The maximum SNR from the ^13^C images of the injected substrate as well as bicarbonate are shown in Fig 5. The SNR of injected lactate was 88 +/-14% of the SNR of injected pyruvate, and the SNR of bicarbonate in the experiments using lactate as the substrate was 52+/-19% of the SNR in the experiments using pyruvate as the substrate.

**Figure 4.**
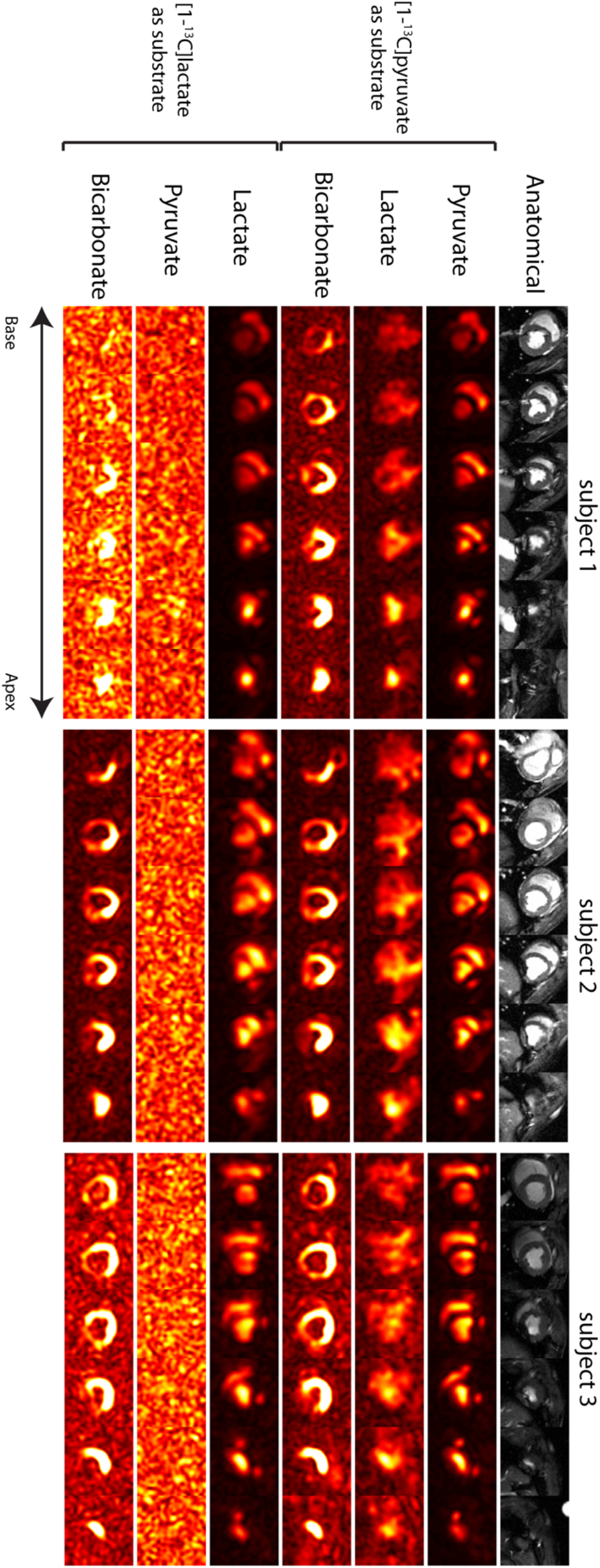
Hyperpolarized 13C imaging data from the hearts of the three animals scanned in this study.

**Figure 5.**
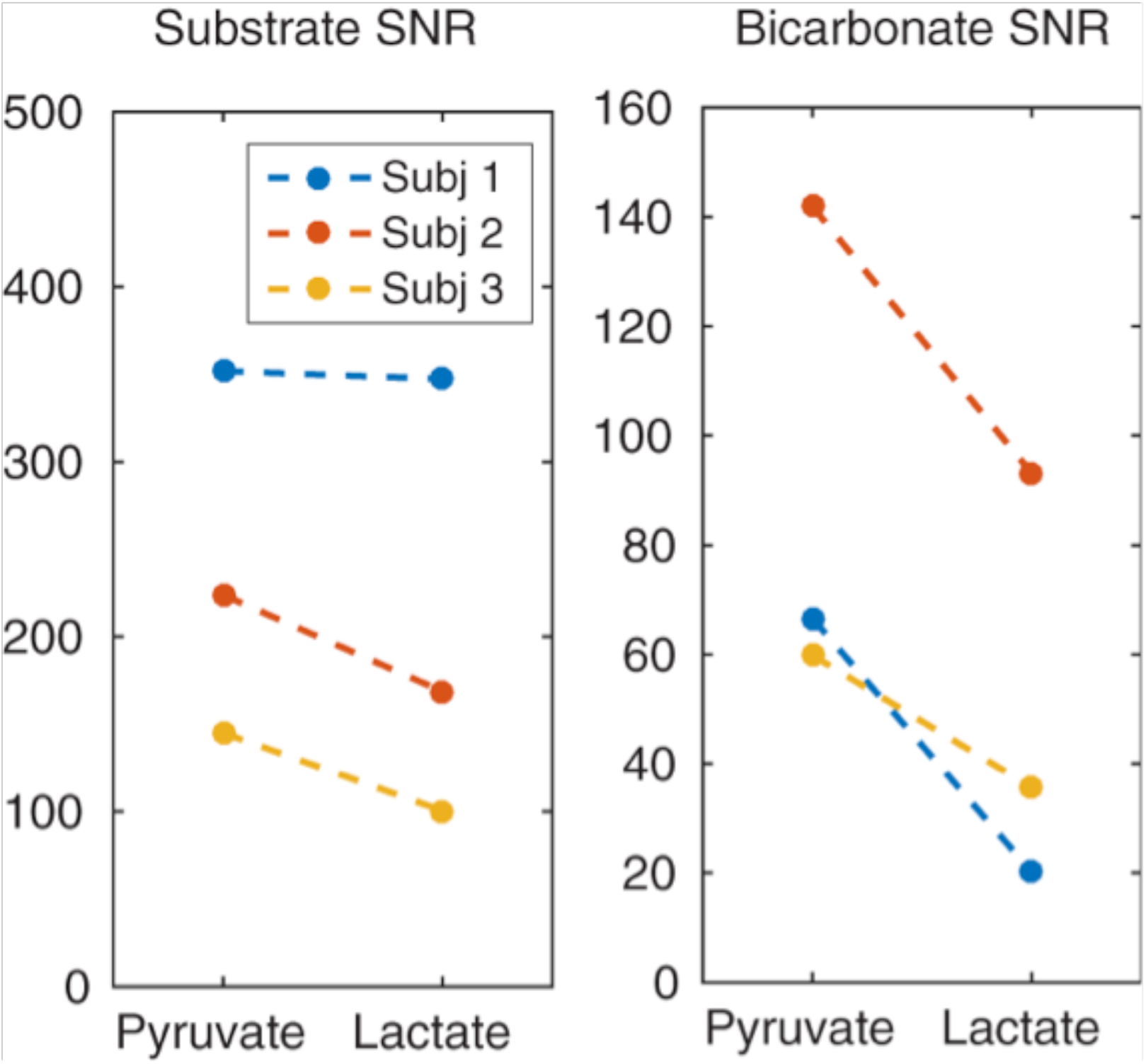
Maximum SNR from the 13C imaging experiments for both injected substrates as well as the metabolic product bicarbonate.

## Discussions

While the safety and feasibility of using hyperpolarized [1-^13^C]pyruvate as a tool to interrogate cellular metabolism has been demonstrated in many studies in humans (2,6,7,13-16), hyperpolarized [1-^13^C]lactate represents an alternative for probing cardiac PDH flux with less perturbation of the metabolic state. At the 0.1 mmol/kg dose that has been utilized in all the human hyperpolarized [1-^13^C]pyruvate studies to date, the injected substrate concentration (∼2 mM after dilution by blood) would greatly exceed its endogenous blood concentration (17). However, using the same dose of lactate results in a similar or lower substrate concentration as compared to its endogenous concentration.

It has also been shown that it is possible to hyperpolarize lactate to a similar level as compared to pyruvate using the Hypersense DNP polarizer system (8-10) and it should also be possible to produce the dose needed for human studies using the Spinlab polarizer (18). However, one potential disadvantage of using hyperpolarized lactate as compared to pyruvate is the shorter T_1_ of lactate and its potential impact on the available magnetization during imaging. Also, to assess changes in cardiac PDH flux using lactate as the injected substrate, the ^13^C labelled molecule passes through two enzyme mediated conversion steps before it is detected as ^13^C bicarbonate, adding a potential mechanism for additional relaxation before the metabolite image is acquired. While prior studies have compared using the two HP substrates in rodent hearts, the *in vivo* SNR of resulting substrate and metabolic product ^13^C images have not been directly compared using the same dose of both substrates.

In this study, using the same polarization protocol and final sample concentration and volume (and similar polarization level achieved in solution) for both substrates, the ^13^C substrate and ^13^C bicarbonate images in pig hearts were acquired and compared following injection of ^13^C lactate and ^13^C pyruvate into the same subject. Not surprisingly, in experiments using HP lactate, the substrate images had lower SNR, which was most likely due to the shorter T1. The ^13^C bicarbonate images also had lower SNR and by a greater degree. Since the substrate image and the ^13^C bicarbonate image were acquired during the same, relatively short imaging window, this additional loss in sensitivity for ^13^C bicarbonate from using ^13^C lactate as the substrate is potentially due to the additional interaction of the ^13^C labelled molecule with the LDH enzyme.

In human studies using HP substrates to date, the time between dissolution and injection has been substantially longer than in pre-clinical studies (∼60 s vs. ∼25 s) due to the extra quality control and time to load the power injector. This could pose a challenge for HP lactate as a substrate for imaging of human heart. Achieving higher polarization and a faster QC process and sample delivery may be necessary before [1-^13^C]lactate can be routinely used for hyperpolarized ^13^C cardiac imaging. However, it may also be possible to infuse a significantly larger dose of lactate, as compared to pyruvate, and to allow a longer imaging window and thus opportunities for signal averaging.

## Conclusions

In this study, the feasibility of obtaining ^13^C bicarbonate images of the heart following injection of hyperpolarized [1-^13^C]lactate was demonstrated in a large animal model, using the same hardware and data acquisition as those used in prior human studies. Hyperpolarized [1-^13^C]lactate was also compared to [1-^13^]pyruvate for cardiac metabolic imaging at the same dose. Lower ^13^C imaging SNR were observed in the experiments using lactate, but the potential of using lactate as a substrate for clinical cardiac metabolic imaging was demonstrated.

## Acknowledgement

The authors thank Canadian Institutes of Health Research (PJT-152928) and Natural Sciences and Engineering Research Council of Canada (RGPIN-2017-06596) for funding support. The authors are also thankful to Jennifer Barry and Melissa Larsen for assistance with the animal preparation.

## Abbreviations used

HP: Hyperpolarized
DNP: Dynamic Nuclear Polarization

